# Activation of cannabinoid receptor CB1 leads to aberrant myelination in development

**DOI:** 10.64898/2025.12.10.693544

**Authors:** Tania G. Miramontes, Kyla R. Hamling, Ryan A. Doan, Saheli Singh, Hannah Y. Collins, Ben Emery, Cody L. Call, Kelly R. Monk

## Abstract

The endocannabinoid system (ECS) has a widespread role in the development and function of the central nervous system (CNS). Cannabinoid receptors like CB1 and CB2 can be activated with exogenous cannabinoids most popularly known as tetrahydrocannabinol (THC) or cannabis and cannabidiol (CBD). The components of the ECS are expressed early in fetal development, and prenatal exposure to cannabis can lead to structural changes in white matter. White matter is composed of neuronal axons ensheathed in myelin, a lipid-rich insulation that facilitates saltatory conduction and maintains axon integrity. In the CNS, myelin is made by specialized glial cells called oligodendrocytes (OLs), which in addition to neurons also express components of the ECS. However, while several studies have focused on how the ECS regulates neuronal development, there is a limited understanding of its impact on OL development or myelin formation.

Therefore, our current study set out to understand how pharmacological activation of the ECS alters OL differentiation and myelin formation *in vivo*. We administered WIN 55,212-2 (WIN 55), a CB1 and CB2 agonist, to larval zebrafish and longitudinally analyzed OL development and myelination *in vivo*. Interestingly, we observed an increase in non-axonal ensheathments in the spinal cord, which appeared to be surrounding neuronal cell bodies. These non-axonal ensheathments were dependent on CB1, as the addition of WIN 55 in a global CB1 mutant prevented this phenotype. Furthermore, this ectopic cell body ensheathment occurred independently from normal myelination processes, as individual OLs did not exhibit changes in the number of myelin sheaths, sheath length, or total myelin output. This study shows that activation of CB receptors *in vivo* leads to increased non-axonal ensheathment without significantly changing OL differentiation or normal myelin formation. Future studies can further investigate the pathways that drive this phenotype to better understand how exogenous cannabinoid activation can regulate the precision of oligodendrocyte ensheathment.

## Introduction

The endocannabinoid system (ECS) is a neuromodulatory system that consists of endogenous cannabinoids (eCBs), enzymes that synthesize and degrade eCBs, and cannabinoid (CB) receptors. CB receptors type 1 and type 2 (CB1 and CB2, respectively) are the best characterized receptors in the ECS. CB1 is one of the most abundant G protein-coupled receptors (GPCRs) in the brain (Howlett & Abood, 2017; Lu & Mackie, 2020), and is enriched pre-synaptically in neurons starting early in fetal development in the central nervous system (CNS) (Fernández-Ruiz et al., 2000; Mato et al., 2003; Wang et al., 2003). In the adult brain, the ECS is well-characterized in modulating neuronal synapses through retrograde signaling that induces synaptic plasticity (Castillo et al., 2012a; Lu & Mackie, 2020). In the developing brain, the ECS also plays an important role in neuronal migration, neurite extension, and axonal patterning (Berghuis et al., 2007; Harkany et al., 2007; Roland et al., 2014).

As both eCBs and exogenous cannabinoids such as tetrahydrocannabinol (THC), the primary psychoactive component of cannabis, bind and activate CB1 and CB2, it is important to understand how ECS activation from exogenous compounds may play a role in developing brains. The prevalence of cannabis use during pregnancy has increased over two-fold in the last two decades to about 8% nationally (Volkow et al., 2019; Young-Wolff et al., 2021). While clinical studies have predominately focused on increased risk of neurocognitive and neuropsychiatric disorders due to prenatal cannabis exposure (Nashed et al., 2021; Wu et al., 2011; Young-Wolff et al., 2021), a recent study showed changes in white matter pathways in children who were exposed to cannabis prenatally (Evanski et al., 2024). In addition, recent single-cell RNA-sequencing data shows expression of CB1, CB2, and several ECS enzymes in glial cell types such as astrocytes, microglia, and oligodendrocyte (OL) lineage cells (Marques et al., 2018; Zhang et al., 2014). Overall, these data have led to further interest in exploring how the ECS affects glial populations and their endogenous role in neural development and function.

One important glial cell type that expresses ECS genes during development are OL lineage cells, which are the myelinating glia of the CNS including their progenitors. Myelin is the lipid-rich, multilayered membrane that wraps around axons to facilitate the rapid transduction of signals, provides metabolic support for neurons (Fünfschilling et al., 2012; Nave, 2010), and maintains axon integrity (Griffiths et al., 1998; Lappe-Siefke et al., 2003). The formation of myelin is dependent on the successful differentiation of OL precursor cells (OPCs), and in humans OPCs begin to arise prenatally and continue to differentiate postnatally (Back et al., 2001; Jakovcevski, 2009; Jakovcevski & Zecevic, 2005). During differentiation, OPCs must identify up to dozens of axon segments in the vicinity, extend processes along their length, and wrap them several times to establish myelin sheaths. *In vitro*, this process can occur cell-intrinsically on any suitable inert structure above a certain size threshold (Bechler et al., 2015; Lee et al., 2012; Lubetzki et al., 1993), while *in vivo* OLs normally avoid myelinating blood vessels, cell bodies, or dendrites, even if they meet the size threshold, as certain signals repel ensheathing processes from stabilizing on these structures (Djannatian et al., 2019; Redmond et al., 2016; Yalçın & Monje, 2021). Additional signals, like growth factors and neuronal activity, modulate OL lineage progression and the extent or axon specificity of myelin formation (Barres & Raff, 1993; Brinkmann et al., 2008; Watkins et al., 2008; Zuchero & Barres, 2013).

The expression of ECS components across the OL lineage indicates that this pathway may have a regulatory role in OL development and myelination. Studies *in vitro* show that activating CB receptors or modulating the concentration of the endocannabinoid, 2-Arachidonoylglycerol (2-AG) can increase OPC proliferation and differentiation (Gomez et al., 2010, 2015; Molina-Holgado et al., 2002). Furthermore, some studies have suggested that exogenous activation of CB receptors *in vivo* induces OL differentiation (Huerga-Gómez et al., 2021) and absence of the CB1 receptor disrupts OL lineage cell progression (Sánchez-de La Torre et al., 2022). This suggests that ECS signaling could provide another mechanism of modulating OPC differentiation and myelin development. However, how exogenous CB activation alters OL development and myelination *in vivo* is not fully understood.

This study aimed to determine how exogenous cannabinoids influence developmental myelination using pharmacological activation and *in vivo* imaging of the larval zebrafish spinal cord at single OL resolution. We show that treating larval zebrafish with a CB1/CB2 agonist increases *mbp:EGFP+* spherical profiles in the spinal cord of the zebrafish by 5 days post-fertilization (dpf), and that this phenotype is mediated by CB1. Additionally, we show that these spherical profiles are non-axonal ensheathments that wrap around neuronal cell bodies without changes to normal OL generation, individual axonal myelination, or axon density. Overall, these data suggests that ECS signaling can impact OL myelin targeting independent of individual OL myelin potential.

## Results

### Exogenous activation of CB receptors increases somatic myelin profiles

Previous studies suggest that activating CB receptors increases OL proliferation and differentiation *in vitro* (Gomez et al., 2011; Ilyasov et al., 2018; Tomas-Roig et al., 2016); however, little is known about how exogenous CB signaling alters OL development and myelination *in vivo.* Therefore, we used stable *Tg(mbp:EGFP-CAAX)* zebrafish to fluorescently label the cell membrane of mature OLs (Fig. 1A) to assess the effects of CB activation on myelination development *in vivo.* We incubated larvae in 0.5 μM or 1.0 μM WIN 55,212-2 (WIN 55; CB1/ CB2 non-selective agonist) or vehicle (1% DMSO) (Hoyberghs et al., 2021) starting at ∼30 hpf, when OPCs are first specified in the zebrafish spinal cord (Park et al., 2002). Larvae were given a fresh dose of WIN 55 along with embryo medium every day until they were imaged at 4 or 5 dpf (Fig. 1A). This continuous WIN 55 treatment had no effect on the gross morphology of zebrafish larvae at 4 or 5 dpf (Supplement 1A). We did not observe differences in mean EGFP+ fluorescence intensity in the dorsal spinal cord across the treatment groups (Supplement 1B, C), suggesting WIN 55 treatment does not drastically alter myelination in the spinal cord. While we did not see evidence of differences in the total amount of myelin, we observed a striking increase in *mbp:EGFP+* spherical profiles, which had a similar size to OL cell bodies, in WIN 55 treated larvae (Fig. 1B, C). While we observed these structures in vehicle-treated controls at 5 dpf (15.2 ± 6.1), WIN 55 treated animals had double the amount (30.4 ± 12.8) at the same age. These data show that WIN 55 does not grossly affect myelination *in vivo* but it appears to alter OL structural features.

**Figure 1:**
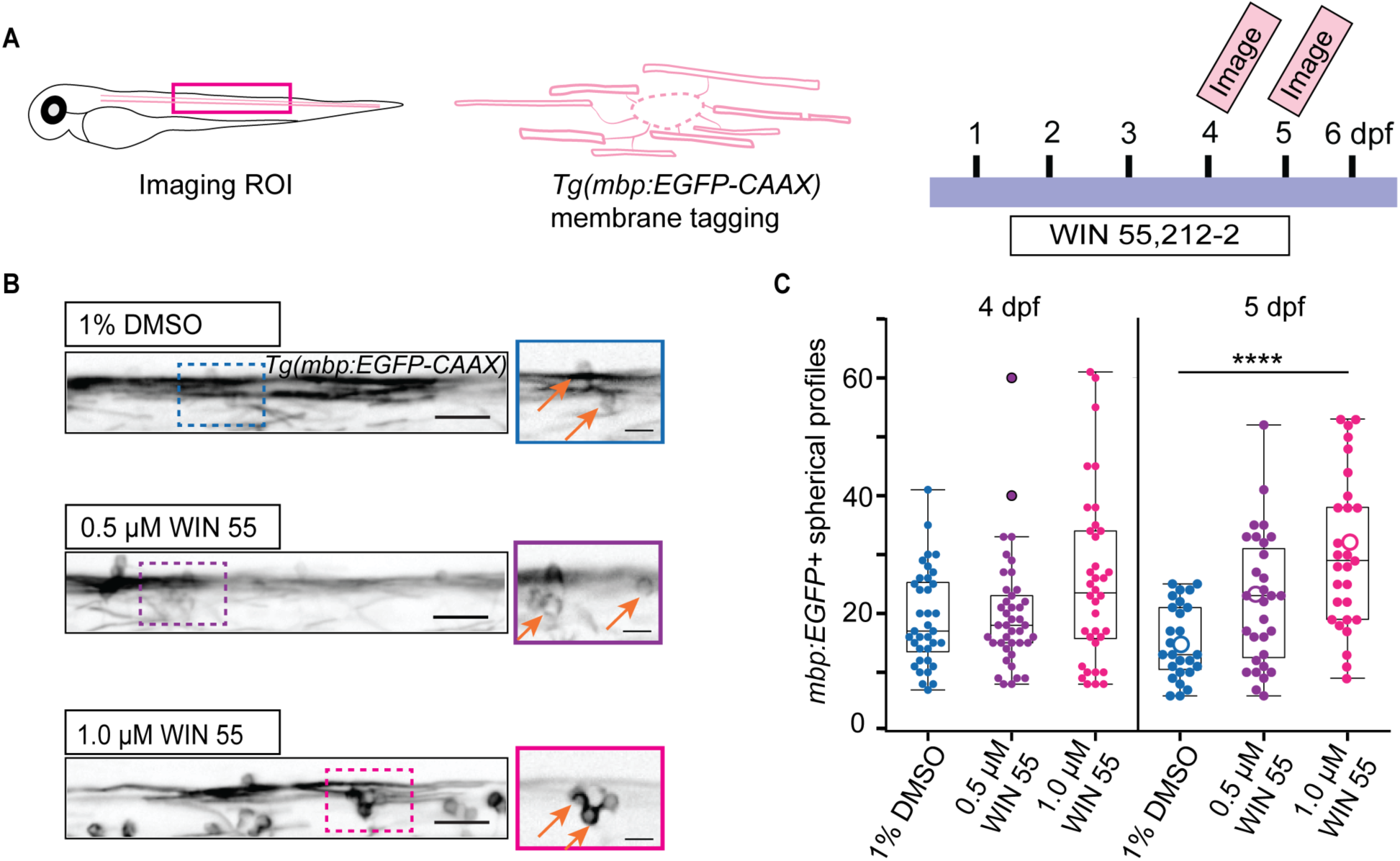
Exogenous application of WIN 55,212-2 during development leads to an increase of *mbp:EGFP*+ spherical profiles. (**A**) Schematic of transgenic zebrafish *Tg(mbp: EGFP-CAAX)* larva highlighting region of interest (ROI; magenta box) and experimental timeline of WIN 55,212-2 (WIN 55) treatment and days imaged (4 or 5 dpf). (**B**) Representative images and enlarged regions of vehicle- and WIN 55-treated spinal cords at 5 dpf. Enlarged regions show *mbp:EGFP+* spherical profiles found in vehicle-treated zebrafish, which are increased in frequency in WIN 55-treated larvae (orange arrows). Left images, scale bar = 25 μm; right images, scale bar = 10 μm. (**C**) Quantification of *mbp:EGFP+* spherical profiles in the dorsal spinal cord at 4 and 5 dpf. Each dot represents one fish. Data depicted as hollow circles correspond to the representative images in (**B**). 4 dpf: DMSO (n = 36), WIN 0.5 μM (n = 41), WIN 1.0 μM (n = 36). 5 dpf DMSO (n = 27), WIN 0.5 (n =31), WIN 1.0 (n = 29). **** p ≤ 0.0001, Kruskal-Wallis test and Dunn’s test with Bonferroni adjustment.

### CB activation causes increased prevalence and stability of non-axonal ensheathments

To further characterize how exogenous CB receptor activation alters OL generation and myelin development we used a double transgenic zebrafish, *Tg(mbp:EGFP-CAAX:mbp:nls-EGFP)* (Fig. 2A), to observe myelinating OL nuclei and OL membrane and distinguish myelinating processes from OL cell bodies.

**Figure 2:**
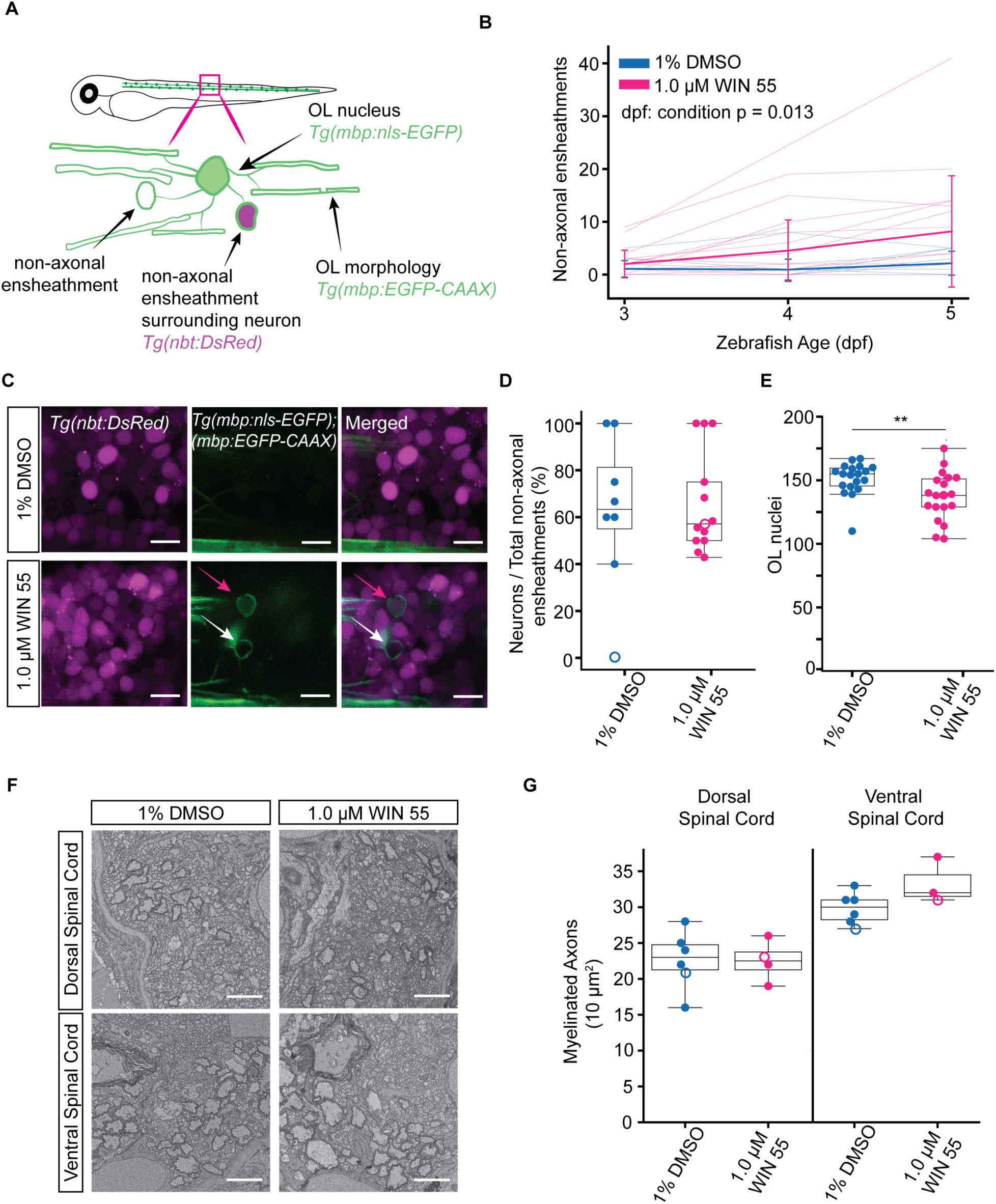
WIN 55 treatment increases neuronal ensheathment over time but does not change the ratio of neuronal ensheathment. (**A**) Schematic of the triple transgenic zebrafish labeling OL nuclei and membrane *Tg(mbp:EGFP-CAAX:mbp:nls-EGFP)* and neurons *Tg(nbt:DsRED)* in the spinal cord. The example OL depicts a non-axonal ensheathment around a neuronal cell body (DsRed filled sphere) or not (unfilled sphere). (**B**) Quantification of non-axonal ensheathments across 3 to 5 dpf. Each thin line represents one fish DMSO (n = 16) WIN 55 (1.0 μM, n = 19). Average values per condition are represented in bold lines graphed with standard deviation. Mixed linear regression dpf: condition p-value = 0.013. (**C**) Representative images of laterally mounted triple transgenic zebrafish. White arrow highlights hollow non-axonal ensheathment and pink arrow shows non-axonal ensheathment around neuron cell body. Scale bar = 10 μm. (**D**) Ratio quantification of how many neurons are ensheathed over total number of non-axonal ensheathments. Two-tailed unpaired t-test, each dot represents one fish. DMSO (n = 8), WIN 1.0 μM (n = 13). (**E**) Quantification of the total OL nuclei in dorsal and ventral spinal cord from a ∼680 μm field of view in zebrafish treated with vehicle and WIN 55 (1.0 μM) at 5 dpf. Each dot represents one fish. DMSO (n = 19), WIN (1.0 μM n = 19). Two-tailed unpaired t-test, ** = p ≤ 0.01. (**F**) Transmission electron micrographs of zebrafish spinal cord cross-sections at 5 dpf after vehicle or WIN 55 (1.0 μM) treatment. Scale bar = 2 μm. (**G**) Quantification of the number of myelinated axons in the dorsal and ventral spinal cord of a 10 μm^2^ region of drug treated fish at 5 dpf. Each dot represents one fish, data depicted as hollow circles correspond to the representative images in (**F**). Dorsal: DMSO (n = 8), WIN 55 1.0 μm (n = 4); Ventral: DMSO (n = 5), 1.0 μm WIN 55 (n = 3). Two-tailed unpaired t-test.

After WIN 55 treatment we observed an increase in hollow EGFP+ spherical profiles (8.2 ± 2.3) compared to vehicle treated larvae (2.1 ± 2.3) by 5 dpf (Fig. 2B), suggesting they were not OL cell bodies but rather non-axonal ensheathments (Fig. 2A). These observed non-axonal ensheathments are commonly observed during normal development when OLs ectopically wrap neuronal cell bodies, but they are typically resolved within the first few days of myelination (Almeida et al., 2018; Djannatian et al., 2019). Therefore, to determine if these non-axonal ensheathments were surrounding neuronal cell bodies, we crossed the double transgenic zebrafish *Tg(mbp:EGFP-CAAX:mbp:nls-EGFP)* zebrafish with a pan-neuronal transgenic animal *Tg(nbt:DsRed)* (Fig. 2A). Of those non-axonal ensheathments we observed, the majority wrapped neuronal cell bodies (Fig. 2C, Supplement Movie 1). While WIN 55 treatment increased the incidence of non-axonal ensheathments, the proportion of neuronal cell bodies wrapped did not change across conditions (71.7% ± 22.1% DMSO, 65.9% ± 21.3% WIN 55) (Fig. 2D). Since not all non-axonal ensheathments surrounded a neuronal cell body, we asked if other cell types like astrocytes were being wrapped. Astrocyte density was unchanged by WIN 55 treatment (Supplement 2 A, B), and we did not observe any non-axonal ensheathments around astrocyte nuclei in vehicle- or WIN 55-treated larvae (Supplement 2 C, D, Supplement Movie 2). Altogether, these results show exogenous cannabinoid exposure during developmental myelination increases non-axonal wrapping of cell bodies but does not alter the specificity of this ectopic ensheathment.

Prior work has established that excessive ectopic cell body myelination can occur when there is an increase in myelinating OLs or a decrease in the number of axons available to myelinate (Almeida et al., 2018; Panlilio et al., 2023). To determine if the increase in non-axonal ensheathments after WIN 55 treatment could be due to an increase of myelinating OLs or a decrease of axons, we quantified the number of mbp:nls-EGFP+ nuclei in both the dorsal and ventral spinal cord. There was a modest decrease in the number of OLs at 5 dpf (Fig. 2E). We then determined if the number of myelinated axons in the spinal cord had decreased, thus reducing the potential area for mature OLs to wrap. Using transmission electron microscopy (TEM) (Fig. 2F), we observed no differences in the number of myelinated axons in either the dorsal or ventral spinal cord (Fig. 2G) at 5 dpf following WIN 55 treatment. Together, these data indicate that the increase in non-axonal ensheathments after WIN 55 treatment is not due to an increase in the number of myelinating OLs nor a decrease in the number of myelinated axons in the spinal cord, suggesting CB activation directly causes OL membrane mistargeting.

### Multi-day CB1 receptor activation is needed to significantly increase the number of non-axonal ensheathments in the spinal cord

The data thus far have shown that exogenous CB stimulation during development leads to increased aberrant myelination of predominately neuronal cell bodies that cannot be explained by changes in OL or axon number. Since WIN 55 is a CB1 and CB2 agonist, we asked if this phenotype was mediated by a single receptor or if activation of both receptors was necessary. In a previous study, we observed upregulation of ECS enzymes in myelinating OLs (Collins et al., 2025), suggesting the ECS is active in these cells. We first focused on CB1 receptor activation, given the single cell sequencing data suggesting higher expression of the gene encoding the CB1 receptor in OL lineage cells compared to CB2 (Marques et al., 2018; Zhang et al., 2014). Therefore, to determine if this phenotype was mediated by CB1 activation, we created a stable mutant of the CB1 receptor gene, *cnr1.* A single guide RNA (sgRNA) was targeted to the 5^th^ transmembrane helix in the second exon of the *cnr1* gene (Fig. 3A), creating a 10 bp insertion and premature stop codon. This mutant allele was then given an allele designation of *vo103* (Fig. 3B; Supplement Fig. 3).

**Figure 3:**
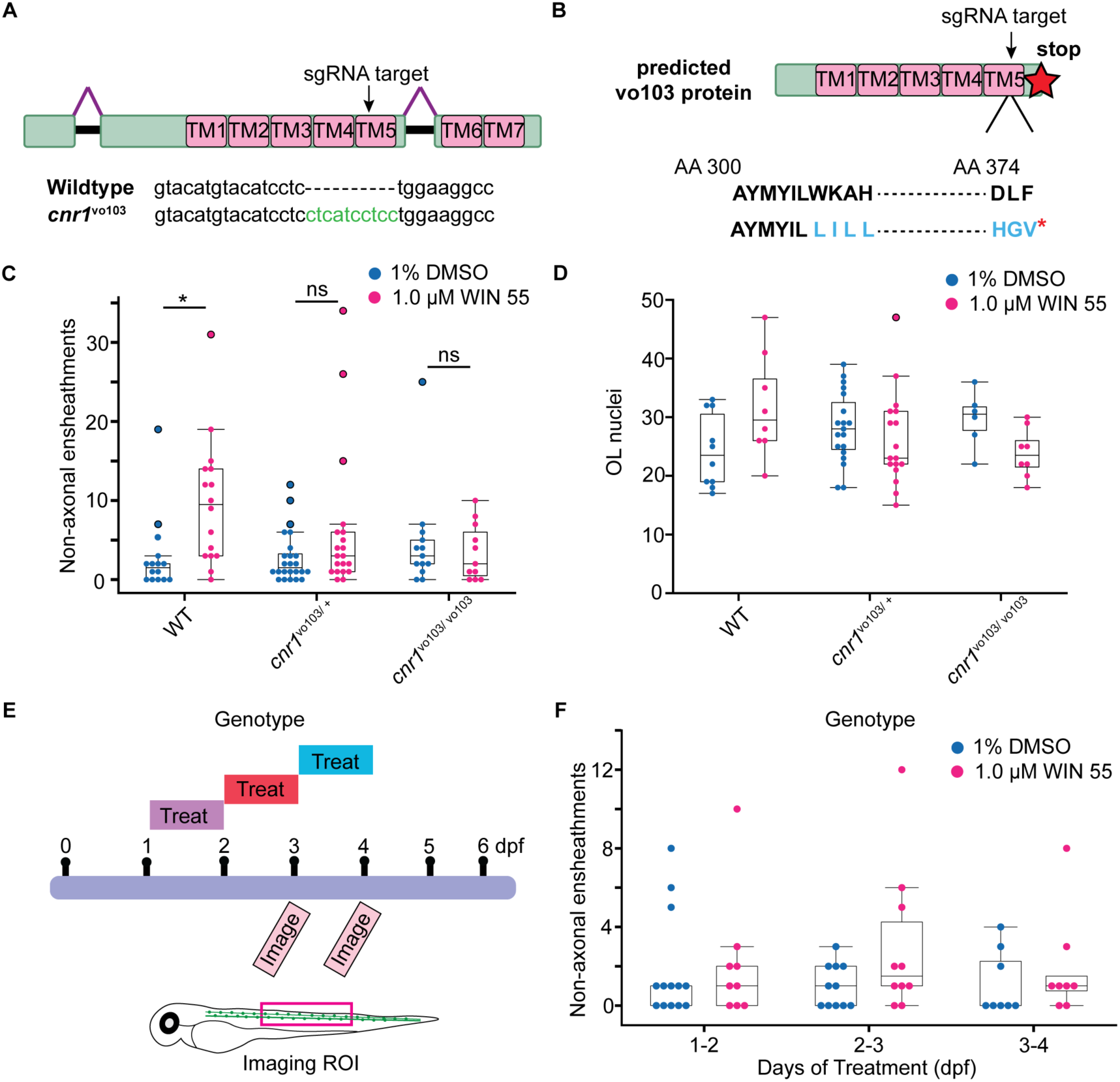
*cnr1* disruption reduces the number of non-axonal ensheathments after WIN 55 treatment. (**A**) Diagram of *cnr1* exons with location of transmembrane helices and the designed sgRNA targeting the end of 5^th^ transmembrane helix. CRISPR-Cas9 mutagenesis using this sgRNA target resulted in a 10 bp insertion. (**B**) Diagram of predicted *cnr1^vo103/vo103^* protein product. 10 bp insertion results in frameshift mutation at amino acid (AA) 306 leading to an early stop codon at AA 376. (**C**) Quantification of the number of non-axonal ensheathments that are not nls-EGFP+ and the (**D**) number of OL nuclei in vehicle- and WIN 55-(1.0 μM) treated fish across wild-type and global heterozygous *cnr1^vo103/+^* and homozygous *cnr1^vo103/vo103^* zebrafish in a ∼ 300 μm field of view. Two-way ANOVA, Tukey’s post-hoc comparison. * = p ≤ 0.05. (**E**) Schematic of experimental timeline. Transgenic zebrafish larvae *Tg(mbp:EGFP -CAAX and mbp:nls-EGFP)* were treated at timepoints either from 1-2 dpf, 2-3 dpf, or 3-4 dpf and imaged at 3 and 4 dpf. (**F**) Quantification of the number of non-axonal ensheathments per treatment condition in vehicle- and WIN 55- (1.0 μM) treated zebrafish imaged at 4 dpf. Two-way ANOVA. DMSO treatment: 1-2 dpf (n = 13), 2-3 dpf (n = 11), 3-4 dpf (n = 8). WIN 55 treatment: 1-2 dpf (n = 9), 2-3 dpf (n = 10), 3-4 (n = 8).

We proceeded to use the double transgenic line *Tg(mbp:EGFP-CAAX:mbp:nls-EGFP)* to quantify the number of non-axonal ensheathments in the stable *cnr1^vo103/vo103^* mutants and their heterozygous and wild-type siblings. We saw no difference in the number of non-axonal ensheathments in the global *cnr1* mutant animals treated with WIN 55 compared to mutant vehicle controls (Fig. 3C). To test whether the absence of a WIN 55 effect on non-axonal ensheathments was due to a change in the number of OLs in the mutant zebrafish, we also quantified the number of non-EGFP+ cells in the dorsal spinal cord across all treatments and genotypes and saw no significant differences (Fig. 3D). These data suggests that the increase in non-axonal ensheathments upon WIN 55 administration is mediated through CB1 receptor activation.

The *cnr1^vo103^*mutants demonstrate that increased aberrant myelination is occurring through CB1 receptor activation. However, the previous experiments were results from a prolonged period (3-4 days) of WIN 55 treatment. Therefore, to determine if both the length of receptor activation and/or a specific OL developmental time point was necessary for the increase of non-axonal ensheathments, we treated zebrafish larvae for a shorter period. We treated the double transgenic zebrafish *Tg(mbp:EGFP-CAAX:mbp:nls-EGFP)* with WIN 55 for a 24-hr period from 1-2, 2-3, or 3-4 dpf, and then imaged at 3 and 4 dpf (Fig. 3E). We assessed how many non-axonal ensheathments were present by 4 dpf and saw no significant changes between vehicle- and WIN 55-treated larvae in any 24-hr treatment period (Fig. 3F). These data suggest that duration of CB receptor activation may be important for causing the increase in mistargeted myelination and that prolonged ECS signaling can alter OL myelin formation.

### WIN treatment does not impair axonal myelination

Our results thus far have shown that CB activation globally increases the number of non-axonal ensheathments, but the above approaches could not distinguish how myelination was affected at the level of individual OLs. To address how CB receptor activation affects the ability of individual OLs to myelinate axons, we sparsely labeled OL lineage cells in *Tg(sox10:KalTA4, UAS:myrEGFP)* zebrafish and measured the number of myelin sheaths, average myelin sheath length, and total myelin output per cell in WIN 55 and vehicle-treated larvae in the same fish across development.

Previous investigations of zebrafish myelin, including from our lab, analyze OL development and myelination based on larval age (Bouchard et al., 2024; Braaker et al., 2025; Choi et al., 2023; Li et al., 2022; Russell et al., 2025). However, this approach cannot accurately distinguish between global delays in myelination versus intrinsic deficits in OL maturation. Additionally, oligodendrogenesis happens across multiple days in the zebrafish and across development in mammals (Ackerman & Monk, 2016; Bartzokis et al., 2012; Matsui et al., 2016; Young et al., 2013). For example, a single timepoint analysis of OL morphology showing a decrease in myelin output can either be due to reduced ability of an individual OL to myelinate, or a global delay in differentiation, effectively creating more immature OLs at later timepoints. We show that in control conditions, the cell age of an OL (days from differentiation) can significantly change analysis of both sheath length and total myelin output, at a single timepoint (5 dpf) (Supplement 4 A-E) and across multiple timepoints (3-6 dpf) during zebrafish development (Supplement 4F-H). Therefore, as an alternative approach, we analyzed the myelinating capacity of individual OLs by grouping OLs based on cell age (days from differentiation) rather than age of the larvae (Fig. 4A). We began imaging larvae at 3 dpf and tracked individual OL lineage cells in the dorsal spinal cord between body segments 5-17 until 6 dpf (Fig. 4 A, B). An OL was considered 1-day-old if at least 80% of the processes appeared to be producing myelin sheaths and it was observed to be an OPC the previous day, regardless of animal age. If an OL cell was first observed with myelin sheaths in a 3 dpf larva, the cell was considered 1-day-old. Interestingly, WIN treatment did not significantly alter the number of myelin sheaths per cell, average myelin sheath length, or total myelin output (Fig. 4 C-E). Altogether, the data suggest that WIN treatment does not alter individual OL myelin potential.

**Figure 4:**
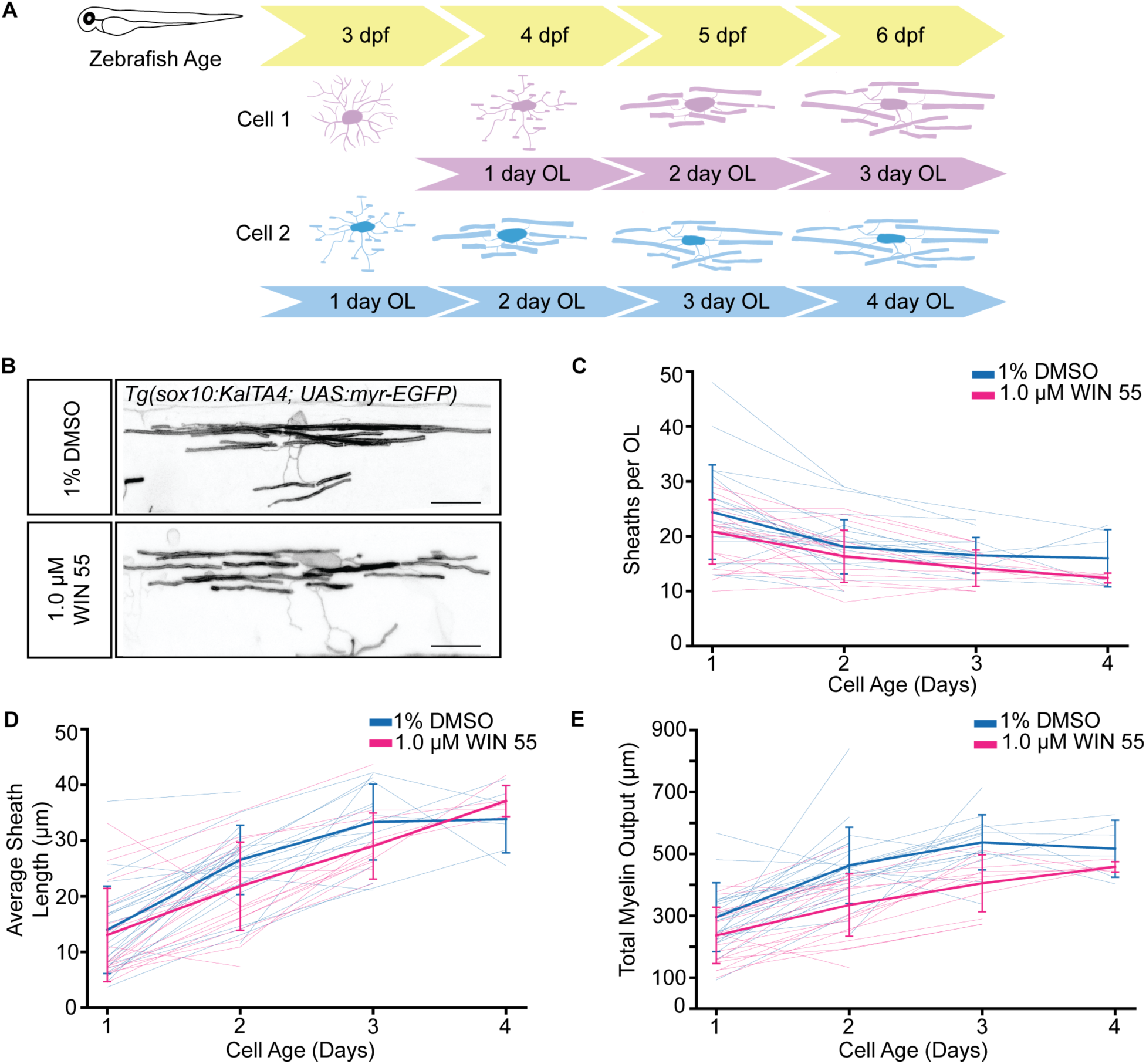
Individual oligodendrocytes treated with WIN 55 develop similarly to control treated oligodendrocytes. (**A**) Schematic of analysis method grouping OL features by cell age compared to standard fish age. (**B**) Representative images of 2-day old OL lineage cells in the dorsal spinal cord of control- and WIN 55-treated fish. Scale bar = 10 μm. (**C-E**) Quantification of (**C**) number of sheaths per OL, (**D**) average sheath length per OL (**E**) total sheath length output in vehicle- and WIN 55- (1.0 μM) treated fish. Each thin line represents single cell data, and the bold lines represent average values with standard deviation. DMSO (n = 26 cells, 13 fish), WIN 55 (1.0 μM, n = 21 cells, 13 fish). Generalized linear mixed effects model.

### WIN treatment allows persistent non-axonal ensheathments across OL development

To further understand how WIN 55 treatment affects individual OL myelin potential, we tracked the presence of non-axonal ensheathments up to 3-day-old OL (Fig. 5 A,B). We then quantified how many non-axonal ensheathments were present per OL and found that vehicle-treated OLs had at most 1 non-axonal ensheathment, whereas WIN 55-treated OLs had up to 3 non-axonal ensheathments (Fig. 5C). Additionally, we found that in vehicle-treated OLs, non-axonal ensheathments were observed the most 1-day post-OL differentiation (4/12 cells) and resolved by the third day (Fig. 5D). However, for WIN 55-treated OLs, the number of OLs with the presence of a non-axonal ensheathment continued to persist and increase (Fig. 5D). Altogether, these data suggest that CB receptor activation leads not only to the increase of non-axonal ensheathments but also their persistence throughout myelin development.

**Figure 5:**
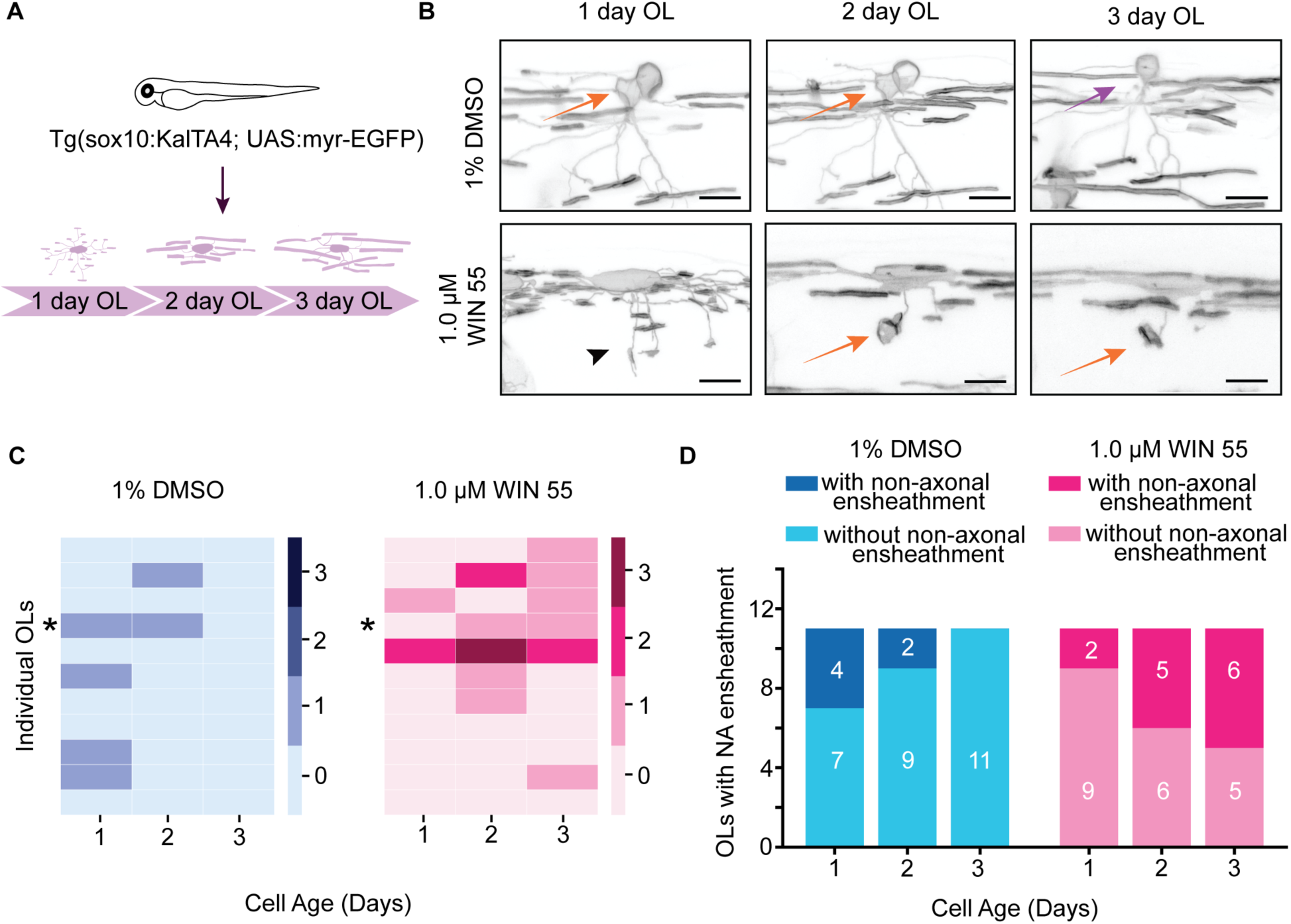
Non-axonal ensheathments fail to resolve in WIN 55 treated OLs and appear at later developmental stages than controls. (**A**) Schematic of transgenic zebrafish used to sparsely label OL lineage cells; each cell that could be tracked for 3 consecutive days was analyzed for the presence of a non-axonal ensheathment process. (**B**) Representative images of individual OLs in vehicle- and WIN 55-treated zebrafish at each day of OL development. Orange arrows highlight presence of a non-axonal ensheathing process, purple arrow highlights where the non-axonal ensheathing process resolved, black arrowhead highlights where a non-axonal ensheathing process will develop. Scale = 10 μm. (**C**) Heatmap depicting number of non-axonal ensheathments observed per OL across cell age in vehicle- and WIN 55-treated zebrafish. Representative images from (**B**) are marked with an asterisk. (**D**) Histogram quantifying how many individual OL treated with DMSO or WIN 55 (1.0 μM) had at least one non-axonal ensheathment forming or formed from the initiation of myelination up to 3 days of OL development.

## Discussion

The roles of the ECS in modulating neuronal activity and neuronal development have been well characterized. While the effects of the ECS in glial cells is beginning to be investigated (Fernández-Ruiz et al., 2000; Stella, 2010), there remains limited understanding of how ECS signaling affects OL development and myelination *in vivo.* As OL lineage cells express several ECS components, the onset of myelination occurs perinatally, and cannabis use during pregnancy is on the rise; understanding how exogenous ECS stimulation impacts developmental myelination is of high importance. Using pharmacological activation of CB receptors while tracking the development of OLs individually and globally *in vivo*, this study reveals that activation of CB receptors caused increased prevalence and stability of non-axonal ensheathments without otherwise impacting normal myelin production. These non-axonal ensheathments are mediated by CB1 receptor activation, helping us further understand the functional impact of ECS signaling on myelin development.

While previous studies of non-axonal ensheathments have shown this phenomenon arises from an imbalance between the myelinating potential of OLs and the abundance of axonal targets (Almeida et al., 2018; Panlilio et al., 2023), our results show that this phenomenon can occur through CB receptor activation independent of these prior reported parameters. Neither OL nor axon densities were changed, suggesting non-axonal ensheathment caused by ECS stimulation is regulated through a distinct mechanism. It has been well characterized how CB receptor activation can modulate neuronal activity (Castillo et al., 2012b; Mackie, 2006; Tseng & Molla, 2025) and it is also increasingly clear that neuronal activity can modulate myelin formation and development (Almeida & Lyons, 2017; Marshall-Phelps & Almeida, 2024; Monje, 2018). Therefore, we hypothesize that activation of CB1 receptors may be altering neuronal activity during development, leading to OLs mistargeting myelin sheaths and that continued exogenous activation is stabilizing these non-axonal ensheathments.

Previous reports have shown that both pharmacological and endogenous activation of CB receptors increases OPC differentiation and myelination *in vitro*. However, our findings show exogenous receptor activation has limited effect of OL lineage progression. Given that multiple components of the ECS are expressed in various cell types in the CNS, it is likely that the synergy of CB activation across cell types *in vivo* have effects unique from CB activation in isolated OPCs. Interestingly, THC administration in postnatal rodents leads to an increase in OPC proliferation and myelination (Huerga-Gómez et al., 2021). While WIN 55 treatment in larval zebrafish did not have the same effect, this difference could be due to the ligand used, as CB receptors are GPCRs and can respond differently to different compounds in different biological contexts. For example, another cannabinoid, Δ^9^-tetrahydrocannabivarin (THCV), which is also present in cannabis can target different receptors in the ECS and have differing effects than THC (Cristino et al., 2020). Our study also focused on the effects of myelination at an individual cellular level, which allowed us to investigate myelin potential through morphological analysis. This study also focused on a narrow developmental window, from early OL lineage specification to the first several days of myelination. Therefore, future work could assess the longer-term effects of CB receptor activation on myelin formation and determine if the non-axonal ensheathments persist throughout myelination or if they eventually resolve.

Given previous literature and our results, future studies can further investigate the extent of various exogenous cannabinoids and their potential effect on OL development and myelin formation. Additionally, while we did not observe changes in axon or astrocyte density in the spinal cord, the question remains as to how signaling on either or both cell types could have altered circuit function. In addition to the evidence showing that CB receptor activation modulates neuronal signaling, it has also been established that astrocytes expressing CB receptors can modulate synaptic transmission (Castillo et al., 2012b; Navarrete & Araque, 2010). Given that changes in neuronal activity can affect OL myelination, it is possible that changes in signaling from all cell types feedback to OL development and exert a unique mechanism leading to mistargeting of myelin and its persistence throughout development. Therefore, future studies can conduct cell-type specific CB receptor knockdown to distinguish the role of various cellular populations on OL development and myelination from exogenous CB activation. Additionally, harnessing the strengths of the larval zebrafish model photopharmacological experiments using inert membrane targeting proteins and photocaged eCBs (Tobias et al., 2021) along with secondary messenger sensors such as cAMP or Ca^2+^ could help us understand the downstream signaling pathways.

Our study provides a better understanding as to how exogenous CB activation affects OL development and myelin formation and provides a new model in which OL can mistarget myelin. Furthermore, this work lays the foundation for understanding how prenatal and perinatal cannabis exposure can affect neurodevelopment.

## Methods

### Zebrafish husbandry

All zebrafish experiments and procedures were done in compliance with the institutional ethical regulations for animal testing and research at Oregon Health & Science University (OHSU; protocol TR02_IP00001148). Zebrafish larvae are fed with rotifers and dry food (Gemma 75) from 5 days post-fertilization (dpf) until 21 dpf. Juvenile fish, (21 dpf until 3 months) are fed using rotifers and dry food (Gemma 150). Adults are fed with a combination of brine shrimp and dry food (Gemma 300). All experiments were approved by the Institutional Animal Care and Use Committee of OHSU.

### Experimental zebrafish

We used the following lines: *Tg(Olig2:dsRed) (*Kim et al., 2008)*, Tg(mbp:EGFP-caax)* (Almeida et al., 2011)*, Tg(mbp:nls-EGFP)* (Karttunen et al., 2017)*, Tg(nbt:dsRed)* (Peri & Nüsslein-Volhard, 2008)*, Tg(10xUAS:myrGFP)* (Li et al., 2022*), Tg(sox10:KalTA4)* (Almeida & Lyons, 2015)*, Tg(glast:myrGFP-P2A-H2AmCherry)* (Chen et al., 2020)*, Tg(glast:myr*GCaMP6s*-P2A-H2AmCherry)* (Chen et al., 2020), nacre pigment mutants, and wild-type AB. All *Tg(mbp:EGFP-caax)* zebrafish were outcrossed for experimental procedures to retain one EGFP copy.

### CRISPR/Cas9-mediated gene disruption and mutant line generation

CHOPCHOP was used to identify a single-guide RNA (sgRNA) to target *cnr1: GTACATGTACATCCTCTGGAAGG* (exon 2) for CRISPR-Cas9 mutagenesis. sgRNA was synthesized using MEGAshortscript T7 Transcription kit (Thermo Fisher). One-cell zygotes were injected with a combination of the sgRNA, 1 Cas9 Nuclease (Integrated DNA Technologies), and phenol red for injection visibility, at a final sgRNA concentration of 25 and 50 ng/nL at a volume between 0.5-1 nL respectively. The cutting efficiency for the guide was verified via enzymatic digestion on the PCR product of the region targeted for editing. Forward and Reverse primers used to test for efficiency and genotyping were Forward-*AACTGCAAGCGGCTCAACTCG* and Reverse- *GACCAGGCTCTTCTGTGACGTG.* Injected F0 larvae were raised to adulthood and outcrossed with wild-type AB zebrafish to establish the F1 generation. Progeny were screened for indels resulting in frameshift mutations using Sanger sequencing. The F1 generation was further outcrossed and used to generate a stable mutant line. For *cnr1^vo103^*larval and adult zebrafish, genotyping tissue digestion was done using Lysis buffer with proteinase-K overnight at 55°C. PCR was performed using GoTaq DNA Polymerase (Promega, M300A) using the same primers stated above. PCR products were digested with Bsl1 (NEB R0555S) at 55 °C for 2 hrs and resolved on a 3% agarose gel with expected mutant band size of 189 bp and band sizes of 130 and 62 bp for wildtype after digestion.

### Mounting fish

For live imaging experiments, larvae were anesthetized with Tricaine (600 μM) in embryo medium and mounted laterally on a dish in 1.2% low-melting-point agarose. Larvae mounted on a dish were pushed to the top of the agarose bead. Agarose was allowed to harden, and the dish was filled with embryo medium containing Tricaine to maintain anesthetized larvae during imaging.

### VAST imaging

We used the VAST BioImager (Union Biometrica) to automate zebrafish imaging from a 96-well plate. One larva was placed per well with 180 μL of embryo medium. Tricane (600 μM) was delivered to the well before larvae were taken up through a capillary system. A water dipping 10×/0.5-NA objective was used with the microscope set up. Microscope specification details can be found in (Early et al., 2018).

### Confocal imaging

All imaging except OL cell counts and EGFP fluorescence measurement was performed on an upright Zeiss LSM 980 confocal with Airyscan 2. A 20×/1.0-NA water dipping objective was used for all LSM 980 imaging, and all images were taken and Airyscan processed using Zen Blue 3.5. We used the following lasers: 488 nm laser (3.0%, 750-800V gain) for EGFP, 561 nm (0.8-1% with 700V gain) for DsRed, 561 nm laser (2.0%, 700V gain) for mCherry.

### Imaging of single cells

For single-cell imaging, Airyscan 2 SR mode was used with a 1.5-4× zoom, depending on the cell size with 0.35-μm z-interval steps. Only cells in the dorsal spinal cord and between body segments 5-18 were imaged and only individual OPCs and OLs that were spatially separated from other labeled cells were chosen for imaging and analysis. The same cells were identified over subsequent imaging days by morphology and body segment from 3 to 6 dpf. If a larva appeared unhealthy (*e.g.*, absent swim bladder inflating or abnormal morphology) it was not imaged for the remainder of the experiment.

### Imaging of non-axonal ensheathments

Larvae positive for both EGFP-CAAX and EGFP-nls expression were mounted laterally and imaged at body segments 7-12. Images were taken using Multiplex SR-4Y mode with a 1.3× zoom and bidirectional scanning at the maximum framerate. The full spinal cord was imaged with 1.0-μm z-interval steps. To determine if non-axonal ensheathments were surrounding a neuron or astrocyte cell body, larvae positive for EGFP-CAAX, EGFP-nls, DsRed, and mCherry expression were mounted laterally and imaged with 0.31-μm z-interval steps.

### Myelinating OL cell counts

To count myelinating OLs, *Tg(mbp:nls-EGFP)* larvae were mounted laterally and imaged using the Zeiss Axio Examiner D1 Spinning disk microscope, 10×/ 0.5 NA objective. All EGFP+ transgenic larvae were imaged with the 488 nm laser (95%, 500 ms exposure time) with 2×2 binning at 1.8-1.84-μm z-interval steps using Zen Blue 2.3.

### Drug Treatment

Zebrafish larvae were enzymatically dechorionated at approximately 24 hpf in bulk with Pronase (1 mg/mL, Millipore Sigma #10165921001). Embryos were incubated and gently swirled for ∼1 min or until the chorion no longer maintained a spherical shape. Embryos were then removed from the Pronase and thoroughly washed with embryo medium 5-6 times before returning to the Petri dish in fresh embryo medium and placed back into the incubator.

(+)- WIN 55,212-2(mesylate) (Caymen Chemicals, 10009023) was dissolved in 100× stock solutions in DMSO and stored at −20 °C. Larvae received fresh embryo medium and WIN 55 daily, therefore a new aliquot was thawed every day and diluted to a 5× concentration using embryo medium. To achieve a final concentration of 1 μM or 0.5 μM WIN 55,212-2 in 1% DMSO, the 5× stock was diluted using embryo medium to the final treatment volume. Control larvae were treated with 1% DMSO. A maximum of 20 larvae were incubated in a total volume of 3 mL per well in a 6-well plate from the beginning of treatment to time of sorting at ∼80-96 hpf. After sorting for fluorescence, larvae were transferred to a 12-well plate and exposed to drug with a total volume of 1 mL drug solution, and a maximum of 6 larvae per well. Larvae were drug-treated starting from ∼30-32 hpf. Every 19-20 h, the larvae were rinsed and left in embryo medium in the incubator for 4-5 h. This was repeated until larvae were imaged. For multiday imaging experiments, larvae were washed with embryo medium after imaging to remove residual Tricaine and retreated with WIN 55,212-2. Concentrations were chosen because they were the highest concentrations that still permitted high viability and normal physical development (Hoyberghs et al., 2021).

### Electron Microscopy

Zebrafish larvae were anesthetized with tricaine and transected between somite 5 and 7 to ensure consistent anterior-posterior sampling. Samples were fixed in modified Karnovsky’s fixative (2% glutaraldehyde, 4% paraformaldehyde in 0.1 M sodium cacodylate buffer, pH 7.4; Electron Microscopy Sciences). Fixation was aided using a PELCO BioWave microwave processor (Ted Pella) with the following cycle: 100 W for 1 min (on/off/on/off), followed by 450 W for 20 s (on/off) repeated five times. Samples were then fixed overnight at 4°C. The following day, samples were rinsed three times (10 min each) in 0.1 M sodium cacodylate buffer. Secondary fixation was performed with 2% osmium tetroxide in 0.1 M sodium cacodylate/0.1 M imidazole buffer (pH 7.5; Electron Microscopy Sciences), using the same microwave cycle as for primary fixation, followed by 2 h incubation at room temperature. Samples were then washed three times (10 min each) in deionized water. En bloc staining was performed with UranyLess (Electron Microscopy Sciences) using microwave assistance (450 W for 1 min, off 1 min, 450 W for 1 min). Samples were washed three times (10 min each) in deionized water. Dehydration was carried out in a graded ethanol series (25%, 50%, 70%, 80%, 95%, 100%). Each step except 70% was microwaved at 250 W for 45 s followed by 10 min at room temperature. Samples were left overnight at 4°C in 70% ethanol. Absolute ethanol steps were microwaved at 250 W (1 min on, 1 min off, 1 min on) with 10 min incubation and repeated three times. Final dehydration used three changes of 100% EM-grade acetone, each microwaved at 250 W (1 min on, 1 min off, 1 min on) followed by 10 min incubation. Infiltration was performed overnight in 1:1 Araldite 812/acetone, followed by two changes of pure Araldite 812. Samples were oriented in flat-embedding molds under a stereomicroscope (Zeiss Stemi 508), allowed to settle for 4–6 h at room temperature, and polymerized at 65°C for ≥48 h. Semithin survey sections (250 nm) were cut, stained with 1% toluidine blue (Fisher Scientific), and examined by light microscopy (Leica DM 300) to verify tissue preservation. Ultrathin sections (70 nm) were collected on Formvar-coated 2×1 mm copper slot grids (Electron Microscopy Sciences), contrasted with UranyLess and 3% lead citrate (Electron Microscopy Sciences), and imaged on an FEI Tecnai T12 transmission electron microscope equipped with an AMT CCD camera.

### Quantification and Statistical Analysis Fluorescence quantification

To analyze EGFP fluorescence, a script was created in Fiji to isolate 400 μm at each z step of the zebrafish dorsal spinal cord. A sum projection image was then created, and the mean fluorescence value was recorded. We quantified myelination in the dorsal spinal cord because it is less dense at that developmental timepoint compared to the ventral spinal cord and allowed for easier quantification.

### OL quantification

The number of EGFP+ cells in *Tg(mbp:nls-EGFP)* zebrafish in both the dorsal and ventral spinal cord were tracked using the multi-point tool on Fiji frame by frame. The entire imaging ROI was used to count the number of EGFP+ nuclei. For Figure 2, the ROI for OL quantification was ∼680 μm. For Figure 3, the ROI for OL quantification was ∼300 μm.

### Quantification of non-axonal ensheathments

An aberrant sphere was identified as an EGFP+ ring viewed first in the EGFP channel. In *Tg(mbp:EGFP-CAAX)* zebrafish, all visible EGFP+ structures were quantified to remove bias and account for possible OL cell body membrane labeling. All EGFP+ sphere structures identified were marked with the Fiji multi-point tool. For larvae with neuronal DsRed expression from *Tg(mbp:EGFP-CAAX:mbp:nls-EGFP)* zebrafish outcrossed with *Tg(nbt:DsRed)*, a composite image was created after marking the non-axonal ensheathments and reviewed frame by frame to determine if the presence of the non-axonal ensheathments appeared within the same frames surrounding DsRed+ cells.

### Single OL analysis

Single OLs were analyzed in Image J’s Neuroanatomy plugin, SNT, to trace individual myelin sheaths. The auto-tracing option was not used during analysis. If two OLs, at relatively the same developmental stage, were in the same ROI and the sheaths were indistinguishable, all myelin sheaths were traced, and the number of sheaths and total myelin output was averaged to represent one cell. All paths were saved as traces and exported to a CSV file, where the number of sheaths, the average sheath length, and total sheath output was calculated per cell. Statistical analyses were performed in MATLAB using the Generalized Linear Mixed Model to determine the effect of drug condition, cell age on sheath length, total myelin output, and average sheath length, considering individual cells and fish as random effects represented as this formula (number of sheaths ∼ condition * cell age + (1|cell) + (1| fish). When analyzing the non-axonal ensheathments from individual OLs we chose to only include cells that were able to be tracked consecutively up to 3 days.

### EM axonal count analysis

Images taken at 2900× magnification were chosen and a 10 x 10 μm square was made and placed in a similar region of the dorsal and ventral spinal cord across fish. Myelinated axons within the 10 x 10 μm square were then counted and analyzed.

### Statistics

All statistical analyses were performed in Python or MATLAB. We tested for normality using Shapiro-Wilk’s test. If data did not pass normality tests, we used Kruskal-Wallis test for data with groups of 3. Animal number, cell number, p-value, are indicated in the figure legends. Animals excluded from the experimental datasets are detailed in the figure legends. All image analysis was done blinded using a Python script to blind filenames. Zebrafish larvae do not have a determined sex during the stages of development employed during the study, so sex cannot be considered a biological variable. For mutant *vo103* experiments, the experimenter was blinded during imaging and analysis on larval genotypes. All representative images in the figures are shown using a unique symbol in the corresponding graphs.

## Code Availability

Custom code and scripts are openly available on GitHub at https://github.com/tmira-018/Figure_Graphs.git and https://github.com/tmira-018/Myelin_analysis_plotly.git or upon request.

## Data Availability

All research materials are available upon request. All data reported in this paper and additional information are available from the corresponding authors upon request. Source data to reanalyze the data is provided in the GitHub https://github.com/tmira-018/Figure_Graphs.git and https://github.com/tmira-018/Myelin_analysis_plotly.git

## Code Availability

Custom code and scripts are openly available on GitHub https://github.com/tmira-018/Figure_Graphs.git. Interactive data visualization for Figure 4 and Figure 5 are also available on GitHub https://github.com/tmira-018/Myelin_analysis_plotly.git

## Acknowledgments

This work was support by NIH/NINDS award F31NS130898 (to T.G.M.), NIH/NIGMS R25GM134978 (S.S.), NIH/NINDS F31NS122433 (H.Y.C.), NIH/NINDS F32NS123005 and K99NS144382 (C.L.C.), NIH/NINDS R01NS138188 (B.E., K.R.M). We thank A. Nechiporuk, J. Frank, J. Williams, J. Li and members of the laboratory of K.R.M. for discussions and critical reading of the manuscript. We thank Suhail Akram, Austin Forbes, Adriana Reyes, and Tia Perry for caring for and maintaining our zebrafish facility as well as the Advanced Light Microscopy Core at OHSU (RRID: SCR_009961).

## Author Contributions

**TGM KRM** conceived the project

**TGM CLC KRM** designed experiments for the project

**HYC** performed foundational proteomics experiments supervised by **BE**

**TGM RAD SS KRH** conducted experiments and collected data for the project

**TGM CLC** analyzed the data and contributed to data analysis

**TGM CLC** wrote the code for image processing and data analysis

**TGM** wrote the manuscript with input from **KRH CLC KRM**

**CLC KRM** supervised the project

---

**Supplemental Figure 1:**
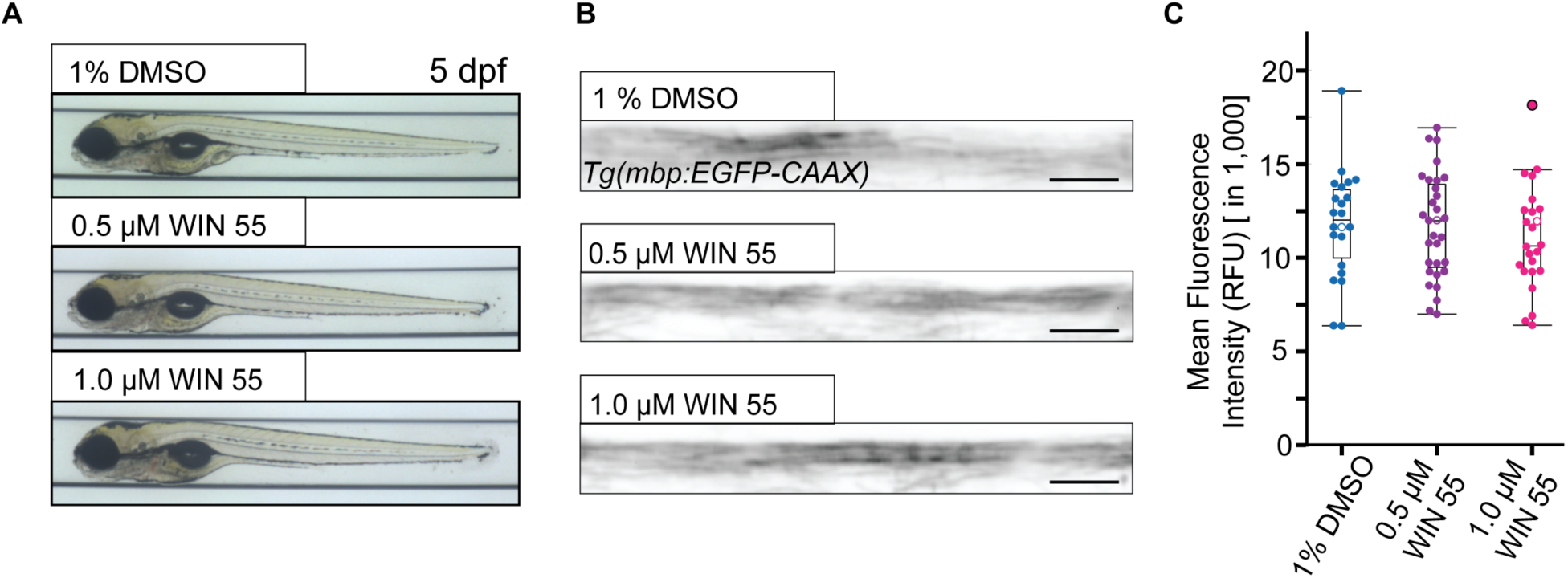
WIN 55 treatment does not change morphological development or overall mbp:EGFP fluorescence intensity. (**A**) Representative images of fish at 5 dpf after vehicle or WIN 55 (0.5 or 1.0 μM) treatment. (**B**) Representative sum projection images of dorsal spinal cord sections used to calculate mean fluorescence intensity values at 5 dpf for vehicle and WIN 55 (0.5 or 1.0 μM) treatment. Scale bar = 25 μm. (**C**) Mean fluorescence intensity quantification of dorsal spinal cord sections at 5 dpf in relative fluorescence units (RFU). Each dot represents one fish, quantification from representative images in (**B**) shown as hollow dots. 4dpf: DMSO (n = 28), WIN 55 0.5 μM (n = 41), WIN 55 1.0 μM (n = 37), 5 dpf: DMSO (n = 22), WIN 55 0.5 μM (n = 31), WIN 55 1.0 μM (n = 24). Kruskal-Wallis Test.

**Supplemental Figure 2:**
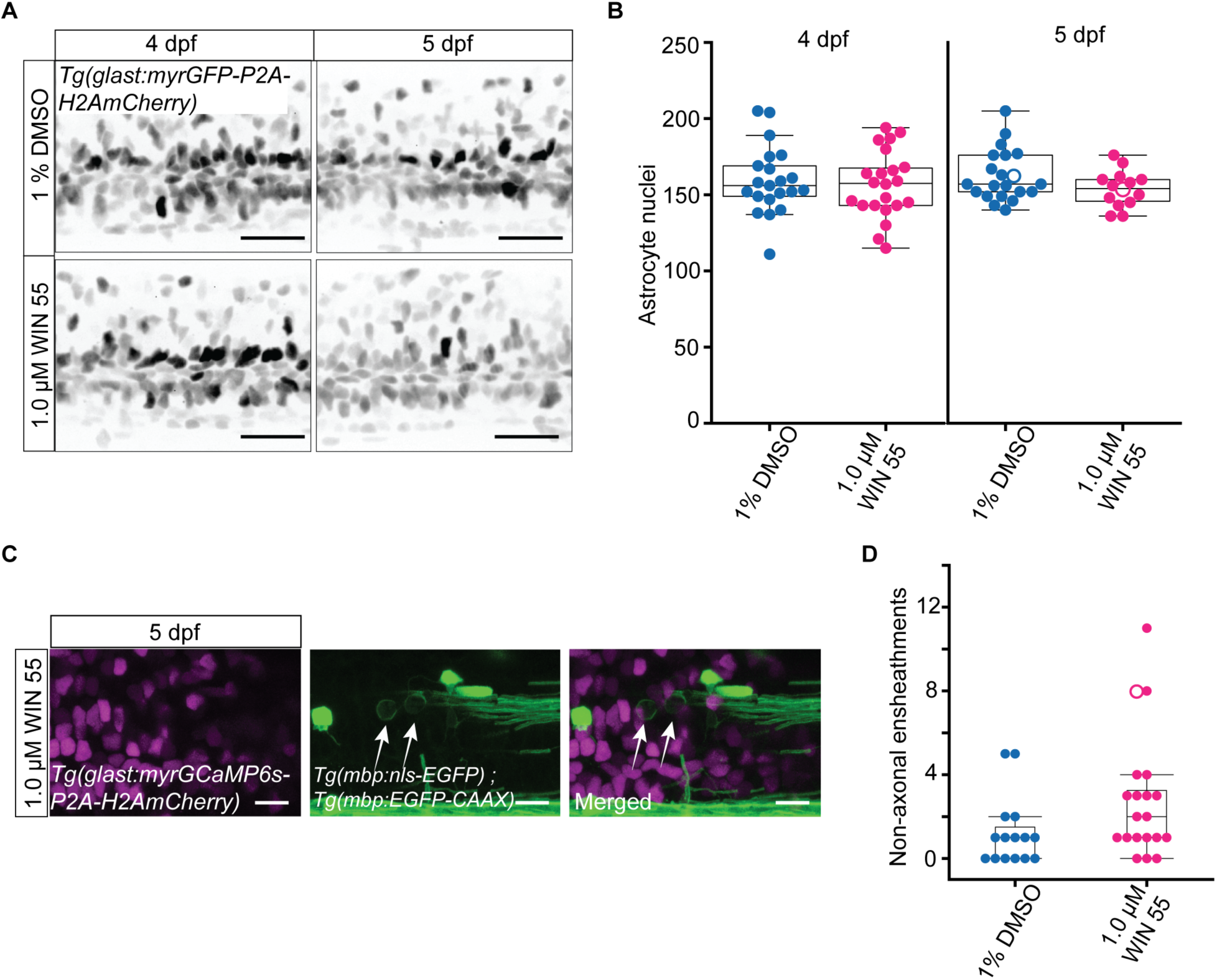
WIN 55 treatment does not affect the number of astrocytes or lead to ectopic wrapping of astrocyte cell bodies. (**A**) Representative max projection images of *Tg(glast:myrGFP-P2A-H2AmCherry)* zebrafish mounted laterally. Zebrafish treated with DMSO control or WIN 55 (1.0 μM) at 4 and 5 dpf. Scale bar = 25 μm. (**B**) Quantification of the number of astrocytes nuclei in the spinal cord of zebrafish at 4 dpf (left) treated with DMSO control (n = 21) or WIN 55 (1.0 μM) (n = 22) and at 5 dpf (right), DMSO control (n = 21) and WIN 55 (1.0 μM) (n = 14). Quantification from representative images in (**A**) shown as hollow dots. Two-tailed unpaired t-test. (**C**) Representative images of laterally mounted transgenic zebrafish *Tg(glast:myrGCaMP6s-P2A-H2AmCherry*) and *Tg(mbp:EGFP-CAAX:mbp:nls-EGFP)* at 5 dpf treated with WIN 55 (1.0 μM). White arrows highlight non-axonal ensheathments. Scale bar = 10 μm. (**D**) Quantification of non-axonal ensheathments in zebrafish treated with DMSO control or WIN 55 (1.0 μM) at 5 dpf. Hollow dot highlights representative image in (**C**).

**Supplemental Figure 3:**
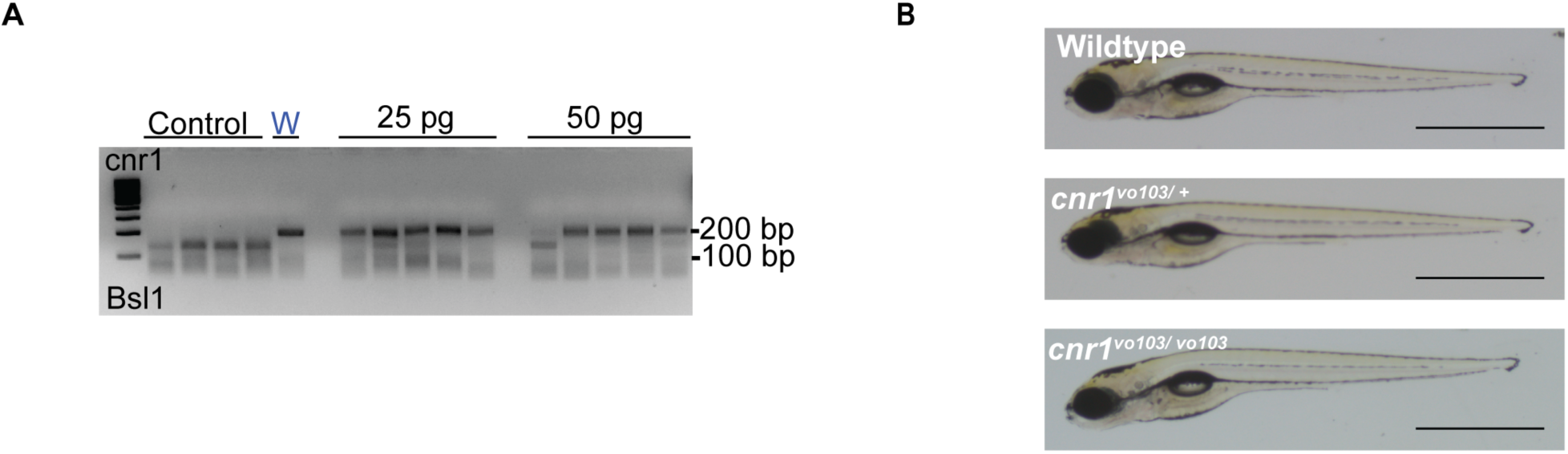
Global *cnr1* mutant zebrafish larvae show no abnormal development. (**A**) Example of amplification of *cnr1^vo103^* allele and restriction enzyme digestion with Bsl1 showing efficiency of Cas9 gene editing at 25 and 50 pg concentrations injected in zebrafish embryos on 3% agarose gel. CRISPR/Cas9 targeting disrupts a Bsl1 cut site resulting in 189 bp band uncut (mutant) and two bands 130 and 62 bp cut (WT). W denotes undigested control. (**B**) Representative images of wildtype, *cnr1^vo103/+^,* and *cnr1^vo103/vo103^* zebrafish larvae at 5 dpf. Scale bar = 1 mm.

**Supplemental Figure 4:**
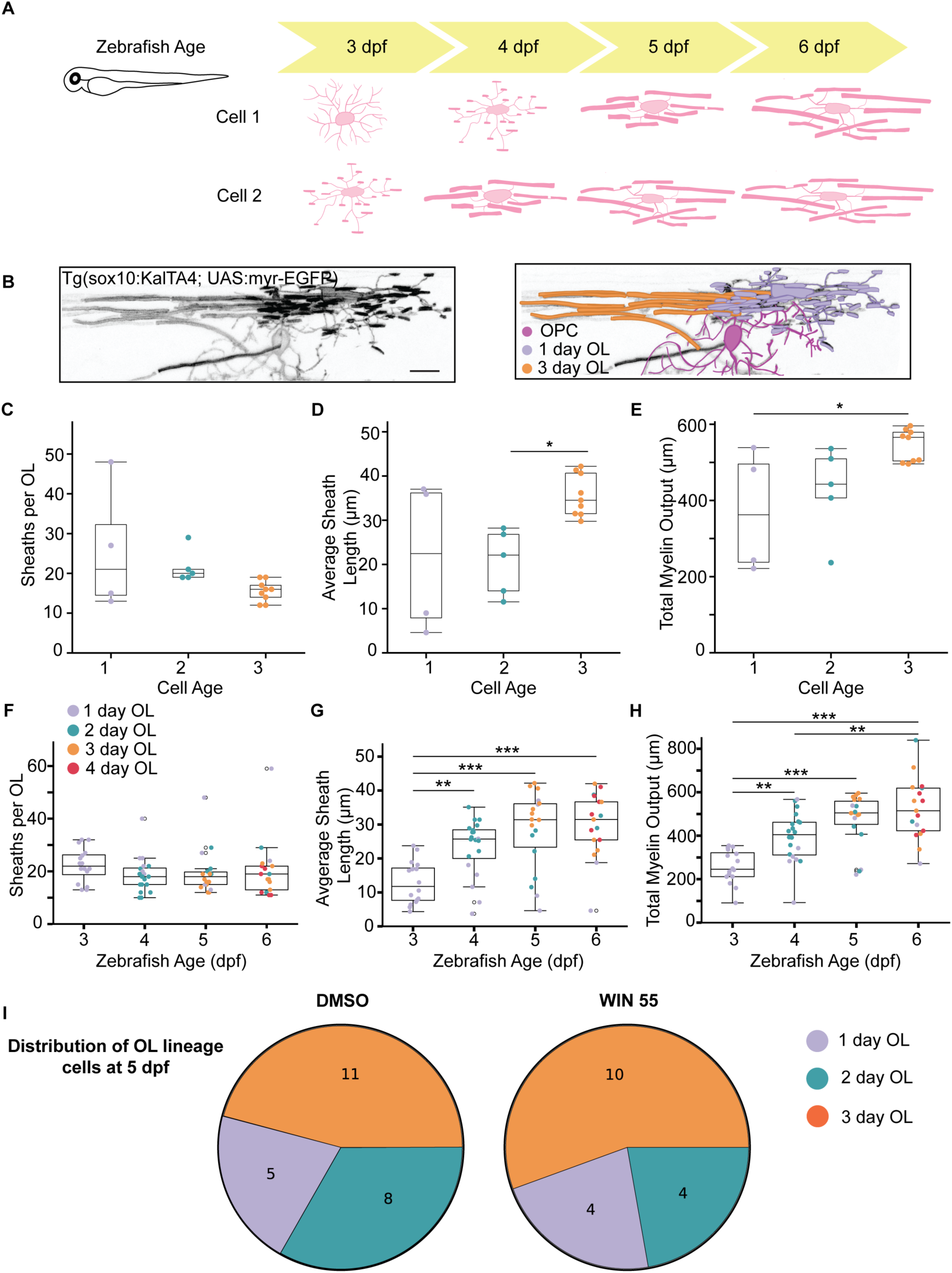
There is significant variability between individual OL cell development in age-matched control animals. (**A**) Schematic demonstrating how the previous method of analysis was matched to the zebrafish age. (**B**) Z at different developmental time points in a zebrafish imaged at 6 dpf. (Left) Max projection image (Right) Pseudo colored image highlighting the 3 different cells. The magenta represents an OPC, the lilac cell represents a cell defined as a 1-day-old OL, the orange represents a 3-day-old OL. Scale bar = 10 μm. (**C - E**) Quantification of the number of cells in each OL cell stage in DMSO treated fish at 5 dpf when quantifying (**C**) the number of sheaths per OL (**D**) Average Sheath Length (**E**) Total Myelin Output. One-way ANOVA, Tukeys HSD post hoc comparison, * = p ≤ 0.05. (**F - H**) Quantification of (**F**) sheaths per OL (**G**) average sheath length (**H**) total myelin output across zebrafish age and the distribution of OL lineage cells of OLs in DMSO control zebrafish larvae at 5 dpf. ** = p ≤ 0.01 *** = p ≤ 0.001. (**I**) Distribution of OL lineage cells in DMSO and WIN 55 treated zebrafish at 5 dpf.

Supplemental Movie 1: Representative video through triple transgenic zebrafish spinal cord Tg(nbt:DsRed) and Tg(mbp:nls-EGFP;mbp:EGFP-CAAX) at 5 dpf, 1% DMSO treated larva on the left side and WIN 55 (1.0 μM) treated larva on the right side. White arrowhead denotes hollow non-axonal ensheathment and pink arrowhead highlights non-axonal ensheathment around a neuronal cell body. Scale bar = 10 μm.

Supplemental Movie 2: Representative video through transgenic zebrafish spinal cord treated with WIN 55 (1.0 μM) *Tg(glast:myrGCaMP6s-P2A-H2AmCherry)* and *Tg(mbp:EGFP-CAAX:mbp:nls-EGFP)* at 5 dpf. Scale bar = 10 μm.

